# Proteome-Wide Analysis of ADAR-mediated Messenger RNA Editing During Fruit Fly Ontogeny

**DOI:** 10.1101/2020.05.07.082404

**Authors:** Anna A. Kliuchnikova, Anton O. Goncharov, Lev I. Levitsky, Mikhail A. Pyatnitskiy, Svetlana E. Novikova, Ksenia G. Kuznetsova, Mark V. Ivanov, Irina Y. Ilina, Tatyana E. Farafonova, Victor G. Zgoda, Mikhail V. Gorshkov, Sergei A. Moshkovskii

## Abstract

Adenosine-to-inosine RNA editing is an enzymatic post-transcriptional modification which modulates immunity and neural transmission in multicellular organisms. Some of its functions are enforced through editing of mRNA codons with the resulting amino acid substitutions. We identified these sites originated from the RNA editing for developmental proteomes of *Drosophila melanogaster* at the protein level using available proteomic data for fifteen stages of fruit fly development from egg to imago and fourteen time points of embryogenesis. In total, 42 sites each belonging to a unique protein were found including four sites related to embryogenesis. The interactome analysis has revealed that most of the edited proteins are associated with synaptic vesicle trafficking and actomyosin organization. Quantitation data analysis suggested the existence of phase-specific RNA editing regulation by yet unknown mechanisms. These results support transcriptome analyses showing that a burst in RNA editing occurs during insect metamorphosis from pupa to imago. Further, targeted proteomics was employed to quantify edited and genomically encoded versions of five proteins in brains of larvae, pupae, and imago insects showing a clear trend towards an increase in editing rate for all of them. Our results may help to reveal the protein functions in physiological effects of RNA editing.

**Significance:** Adenosine-to-inosine RNA editing has multiple effects on body functions in many multicellular organisms from insects and molluscs to humans. Recent studies show that at least some of these effects are mediated by changes in protein sequences due to editing of codons in mRNA. However, it is not known how exactly the edited proteins can participate in RNA editing-mediated pathways. Moreover, most studies of edited proteins are based on the deduction of protein sequence changes from analysis of transcriptome without measurements of proteins themselves. Earlier, we explored for the first time the edited proteins of *Drosophila melanogaster* proteome. In this work, we continued the proteome-wide analysis of RNA editome using shotgun proteomic data of ontogeny phases of this model insect. It was found that non-synonymous RNA editing, which led to translation of changed proteins, is specific to the life cycle phase. Identification of tryptic peptides containing edited protein sites provides a basis for further direct and quantitative analysis of their editing rate by targeted proteomics. The latter was demonstrated in this study by multiple reaction monitoring experiments which were used to observe the dynamics of editing in selected brain proteins during developmental phases of fruit fly.

**Highlights:**

- Proteogenomic approach was applied to shotgun proteomics data of fruit fly ontogeny for identification of proteoforms originating from adenosine-to-inosine RNA editing.
- Edited proteins identified at all life cycle stages are enriched in annotated protein-protein interactions at statistically significant level with many of them associated with actomyosin and synaptic vesicle functions.
- Proteome-wide RNA editing event profiles were found specific to life cycle phase and independent of the protein abundances.
- A majority of RNA editing events at the protein level was observed after metamorphosis in late pupae to adult insects, which was consistent with transcriptome data.
- Targeted proteomic analysis of five selected edited sites and their genomic counterparts in brains for three phases of the fruit fly life cycle have demonstrated a clear increase in editing rate of up to 80% for the endophilin A protein in adult flies.

## Introduction

Editing of a ribonucleic acid by RNA-dependent adenosine deaminases (ADAR) is an ancient mechanism of RNA post-transcriptional modification which occurs in cells of many multicellular eukaryotes [1]. A distinctive feature of enzymatic reactions catalyzed by proteins of this family is that they deaminate adenosine residues in double-stranded RNA (dsRNA) secondary structures [2]. The resulting inosine residues, in contrast to original adenosines, are less affine to the uridine residues and prefer to bind cytidines by hydrogen bonding. Thus, the dsRNA regions become disrupted. The original role of ADAR activity, according to the theory accepted in many studies, is to participate in the non-specific immunity mechanisms [3,4]. For example, in mammalian cells, an excess of dsRNA of any origin causes a specific response initiated by the dsRNA sensors and followed by activation of type I interferon cascade [5]. In case of viral RNA, these responses are necessary, while ADAR enzymes can diminish them, thus acting as a negative feedback and avoiding hyperactivation of non-specific immunity against own dsRNA [6].

Targeting dsRNA by ADAR was shown to be mostly dependent on the secondary structure and also on sequence context, at least for human ADAR1 [7]. If the corresponding secondary structure is situated in exons, the enzyme also deaminates adenosines in its codons. As the resulting inosine residues prefer cytidines to uridines, the amino acid content of encoded proteins may be changed by single amino acid substitutions. Moreover, splice sites subjected to RNA editing, which is shown to occur before mRNA splicing, can influence the protein sequence more dramatically [8]. However, it may be hypothesized that these proteome-associated consequences represented originally just adverse effects of dsRNA deactivation, often non-adaptive and harmful. On the contrary, those non-synonymous editing sites, which are found to be edited recurrently, are obviously under positive selection. That is why natural selection acts differently on the coding and non-coding RNA sites [9]. Moreover, it was shown that the coding mRNA sites, in many cases, are edited at higher rates than the non-coding ones, as the resulting proteoforms acquire a specific role during lifespan of an animal [10].

The role of RNA editing of coding sequences may differ in mammals and insects. In humans and mice, two active ADAR isoforms are found. Of them, ADAR1, which is encoded by *adar* gene, is responsible for deamination of massive dsRNA arrays, e.g. in response to interferon signaling, while ADAR2 (*adarb1*) is thought to target protein-coding transcripts [11]. In fruit flies, a single gene, *Adar*, encodes the deaminase, which is similar to mammalian ADAR2 [12]. However, a recent study focused on functional characteristics of mutated strains of *Drosophila melanogaster* has illustrated that the fruit fly’s enzyme shared functionality of both human isoforms, regulating the dsRNA immune responses and editing the mRNA transcripts [13]. Generally, in fruit flies, coding editing events occur at higher rates than in humans and mice, which was shown at both transcriptome [14,15] and proteome [16,17] levels. This fact, at least partially, can be explained by the need for profound morphological and physiological changes in the insect which experiences a metamorphosis during its lifespan. In *Drosophila*, RNA editing reportedly contributes to maintenance of genome compactness, which is important due to the need of metamorphosis. Edited protein variants provide modified functions that are required at different life phases of the insect [18]

Functional role of editing in proteins from any species is clarified for a small cohort of well-characterized sites. For instance, only a single coding Gln-to-Arg editing site in *gria2* AMPA glutamate receptor subunit is essential for survival of mice. Mimicking this polymorphism by genomic mutations in combination with ADAR inactivation leads to a normal survival of transgene mice without obvious pathology [19]. However, some important functions are attributed to the nonsynonymous RNA editing of filamin A in murine vasculature [20] and coatomer alpha in human cancers [21]. Also, many of known nonsynonymous RNA editing sites, e.g., from fruit fly, are still not characterized functionally.

Coding RNA editing events in mRNA were mostly studied in transcripts [22], and only a few works used mass-spectrometry data to identify amino acid substitutions generated by transcriptome-wide ADAR activity. This was studied for human cancers [21], murine and human brains [17], as well as fruit fly’s body, head and brain [16].

A useful approach to consider a biological trait or process is answering the four questions suggested by the ethologist Niko Tinbergen in his 1963 seminal work [23]. These questions relate to ontogeny, phylogeny, mechanism, and adaptive significance of the trait [24]. In the present work, we aim to describe such a complex trait as a proteome-wide consequence of adenosine-to-inosine RNA editing for *D. melanogaster*’s ontogeny.

It was reported before that the editing of RNA transcripts is differential during ontogeny of various organisms, from insects to mammals. What factors can regulate this differential editing? Normally, enzymatically driven processes are dependent on the enzyme expression. However, it was not confirmed for the ADAR in humans [25]. Further, some of the target-specific RNA binding proteins were found to modulate the enzymatic activity for one mammalian isoform, ADAR2, which was thought to be responsible for differential editing of mRNA targets [26]. For example, SRSF9 is an RNA splicing factor shown to inhibit editing of CYFIP2 mRNA target in human cells [26].

In fruit fly, the *Adar* gene is more expressed in the central neural system, which is similar to the mammalian ADAR2/*adarb1* [27]. Generally, when mutated, it results in deficiencies in locomotive ability and faster neurodegeneration with aging [27]. These effects were recently reversed by autophagy activation, which pointed to impaired proteostasis and increased TOR signaling as important mechanistic clues to explain effects of *Adar* deficiency [28]. Further, its gene expression is boosted at insect metamorphosis [29]. For *Adar* transcription, a complex regulation is described with many known alternatively started and spliced transcripts which are differential in embryos and adult insects [29]. Briefly, a constitutive 4A promotor generates embryonic transcripts, some of which are also characterized by inclusion of additional 3a exon, which encodes a 38-amino-acid protein fragment in a specific 3a ADAR isoform. The 4B promoter is thought to provide exclusion of the 3a exon with generation of adult-specific 3/4 protein isoforms which predominate in the insect after metamorphosis [30]. In adult-specific transcripts, auto-editing was also described, which converts a serine residue to glycine in the resultant proteins. In adult insects, both edited and unedited enzyme isoforms are found [30]. From these data, it can be hypothesized that the transcript diversity during fruit fly ontogeny may provide differential editing of selected targets. However, a detailed mechanism of this process is still to be elucidated.

Recently, we identified the consequences of mRNA editing in adult fruit fly proteomes, by the search in shotgun mass-spectrometry data, specifically, for the brain proteome [16]. The next step was to estimate the dynamics of the ADAR-mediated editome at the protein level, which was done in this work using publicly available deep proteome data obtained for sequential developmental stages of *D. melanogaster* [31]. The data were searched against the genomic database supplemented by possible amino acid substitutions introduced by ADAR editing as described elsewhere [16]. The bioinformatic pipeline was supplemented by experiments, in which both edited and unedited sites in selected proteins were quantified at different developmental phases of the fruit fly brain using multiple reaction monitoring (MRM).

## Results and discussion

### Reanalysis of proteomes of fruit fly ontogeny stages to identify ADAR-edited protein sites

After we started searching for traces of RNA editing in the fruit fly brain proteome [16,32], a comprehensive developmental proteome map of *D. melanogaster* was released [31]. The map was based on a shotgun proteomic approach compatible with the proteogenomic pipeline proposed earlier [16]. Thus, it was tempting for us to reanalyze these new high-quality proteomic data by addition of edited amino acid sequences to the conventional genomic database. As a result, we were able to augment the developmental dynamics of the proteome itself described in the above work [31] with the knowledge on the dynamics of proteins changed by adenosine deamination of their transcripts, the process known to be developmentally regulated [22].

Search for single amino acid polymorphisms, including those generated by RNA editing, is based on peptide spectrum matches (PSMs) from shotgun proteomic data [33]. Proper validation of associated findings is needed to avoid reporting false positive hits, which is especially difficult when the shotgun data are reanalyzed without access to original specimens, such as in the current study. We applied a conservative filtering procedure described previously [16] to the output of the search engine to get rid of dubious identifications. In addition, all hits with chromatographic retention time deviating strongly from theoretically predicted values were also excluded to avoid further doubt due to possibly false positive results [34].

Data to be reanalyzed consisted of two subsets, the first one including fifteen stages of fruit fly development from egg to imago and the second one spanning fourteen time points of embryogenesis, i.e. the egg development [31]. Identified ADAR editing sites filtered as described are listed in Table 1. In total, 42 edited sites were found, each belonging to a unique protein. Of these, only twelve sites were identified in the embryonic dataset, with eight of them overlapping with the life cycle dataset containing also four time points of egg development. The list of edited protein sites was compared to the recently obtained proteomes of whole bodies, heads and brains of adult insects [16]. About one third of sites observed in the current study was also identified in [16] (Table 1). The difference between the identifications can be explained by the presence of phase-specific edited sites which were omitted in the studies on adult animals.

**Table 1.**
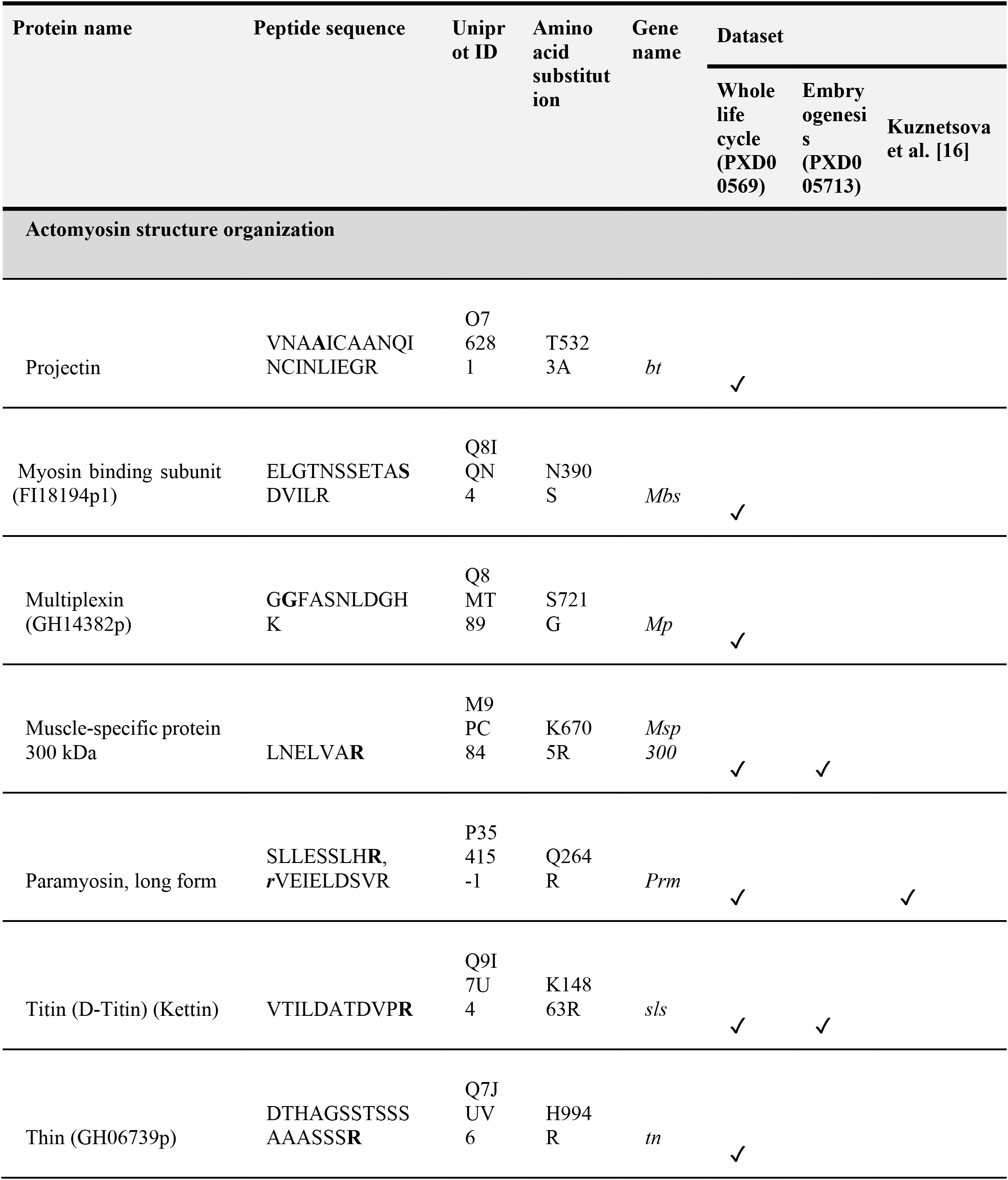
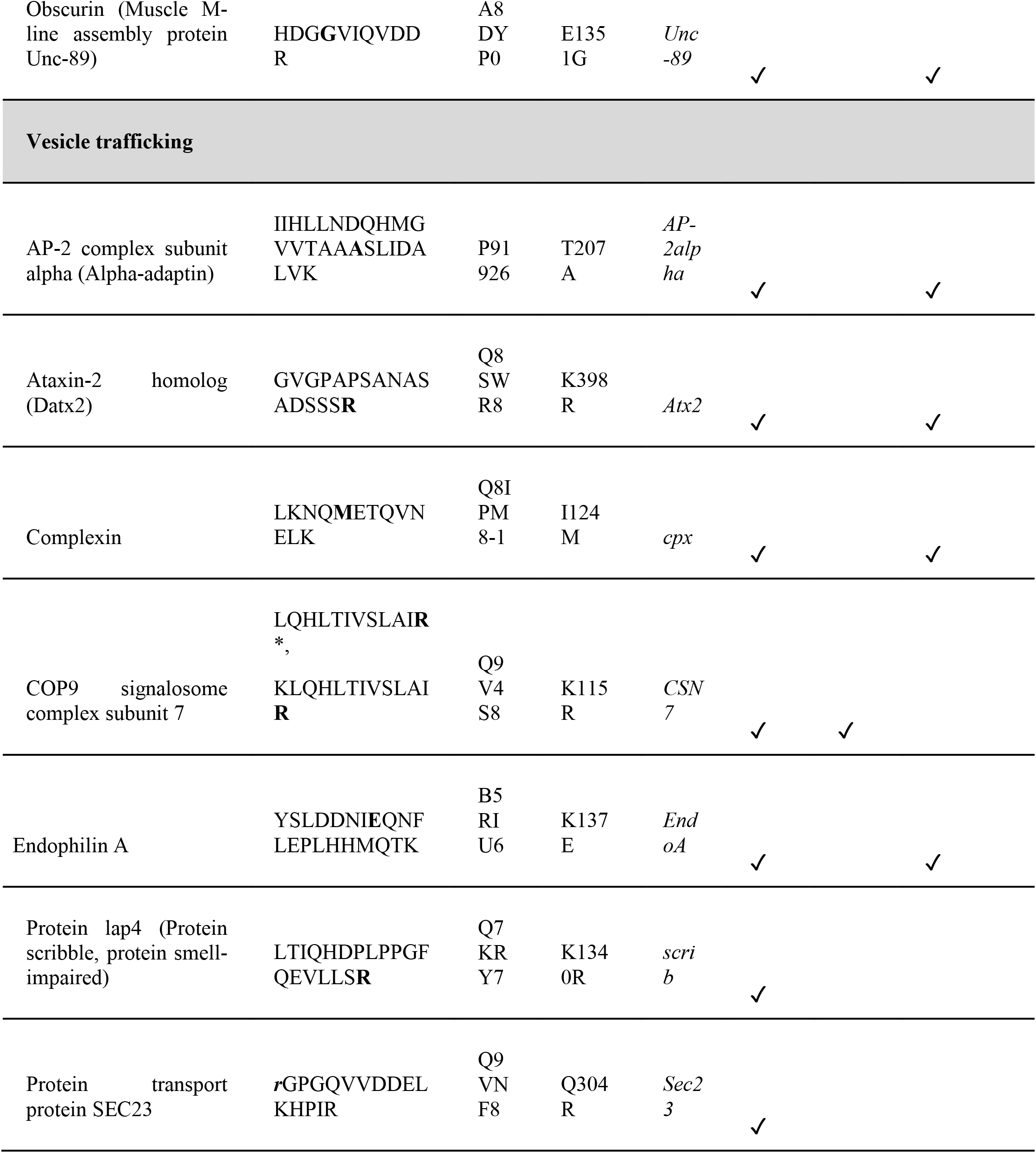
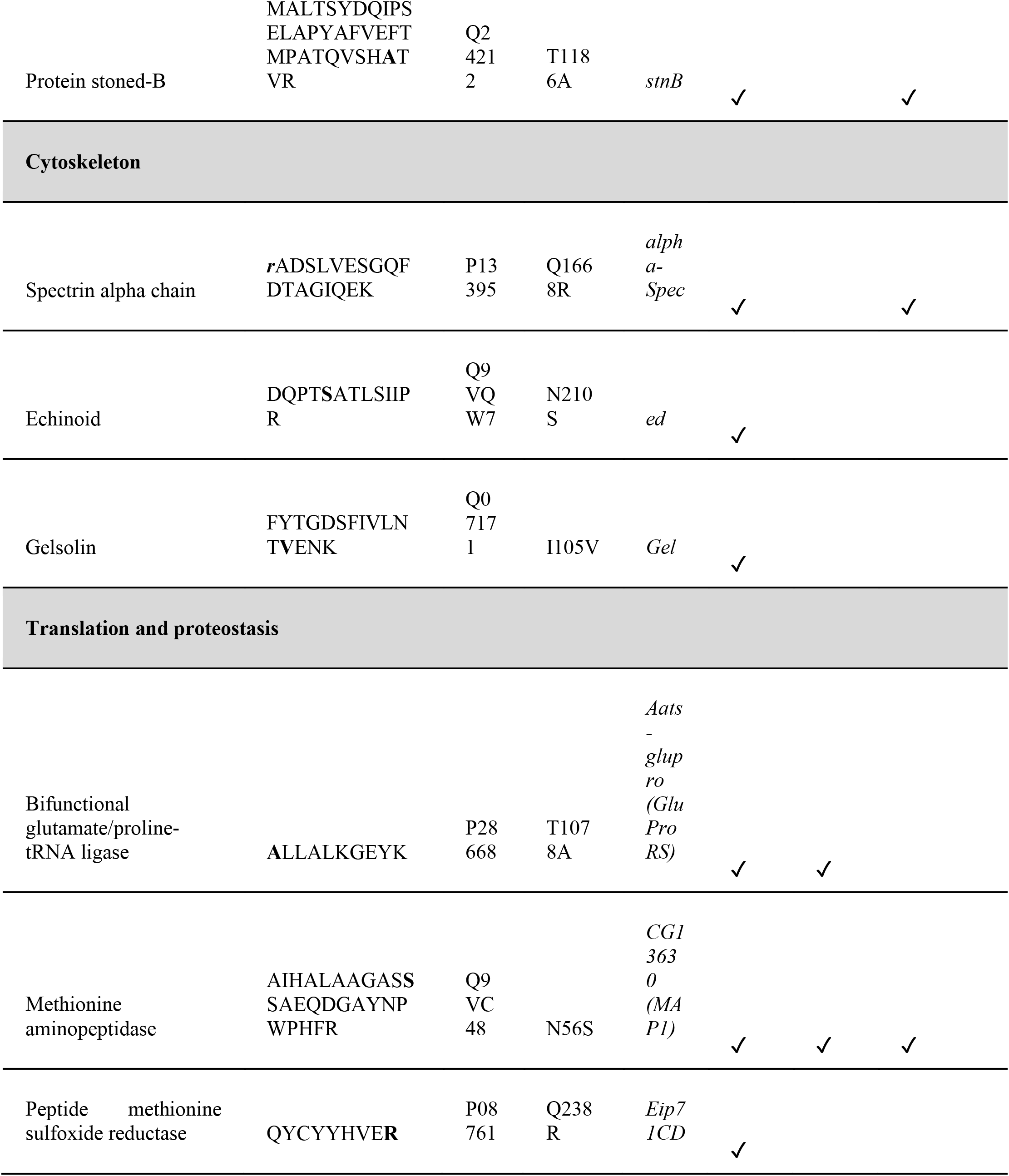
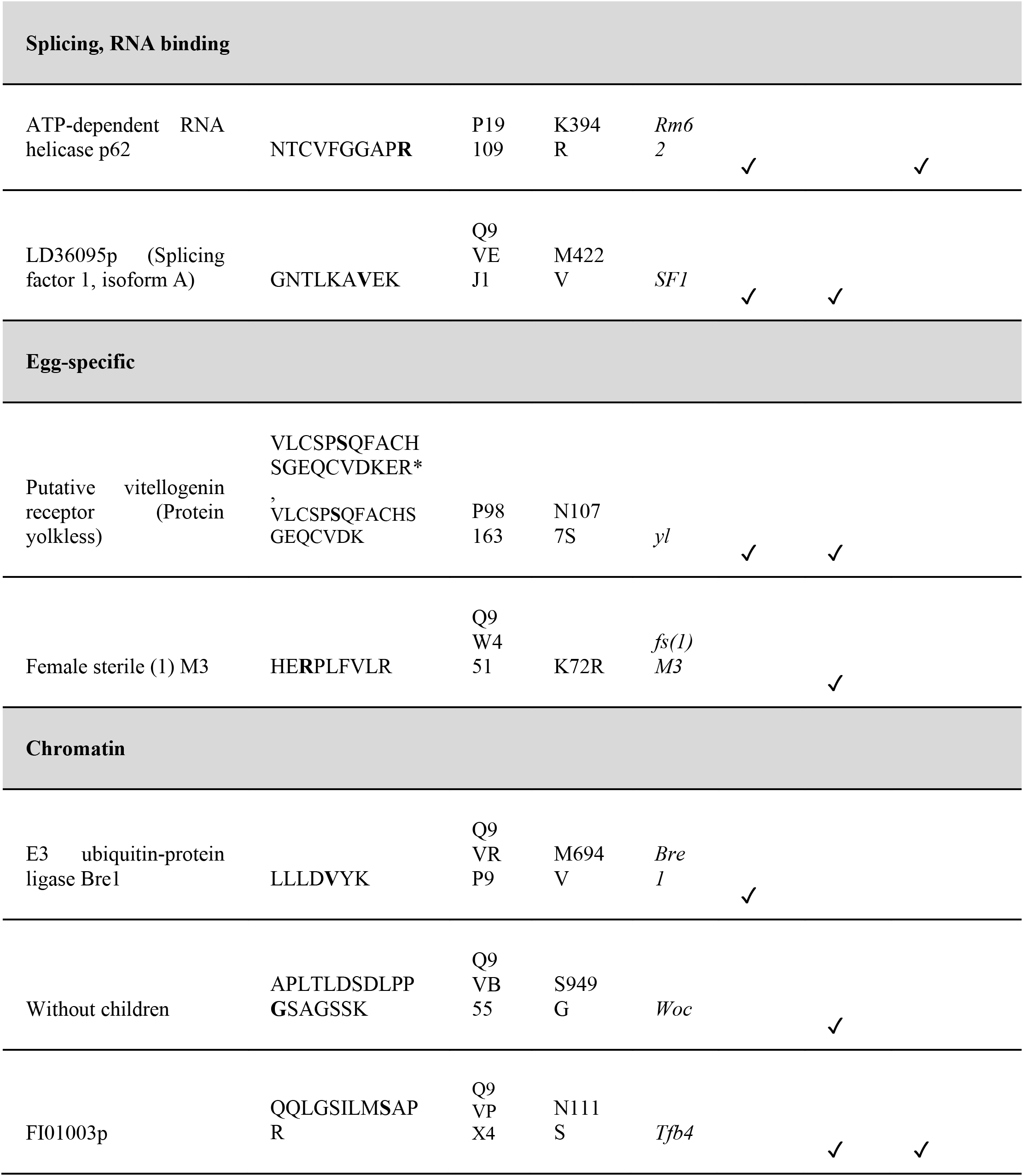
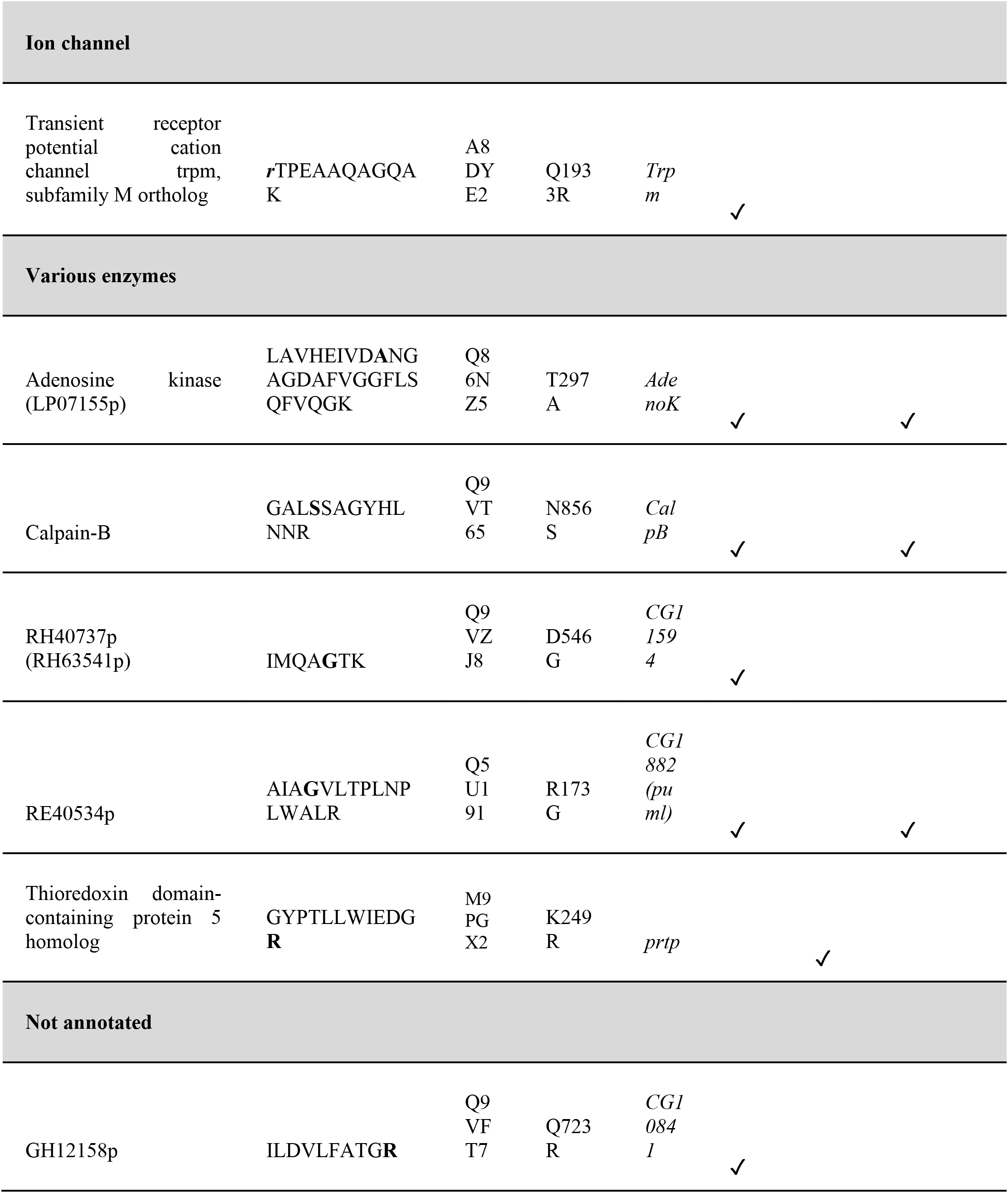
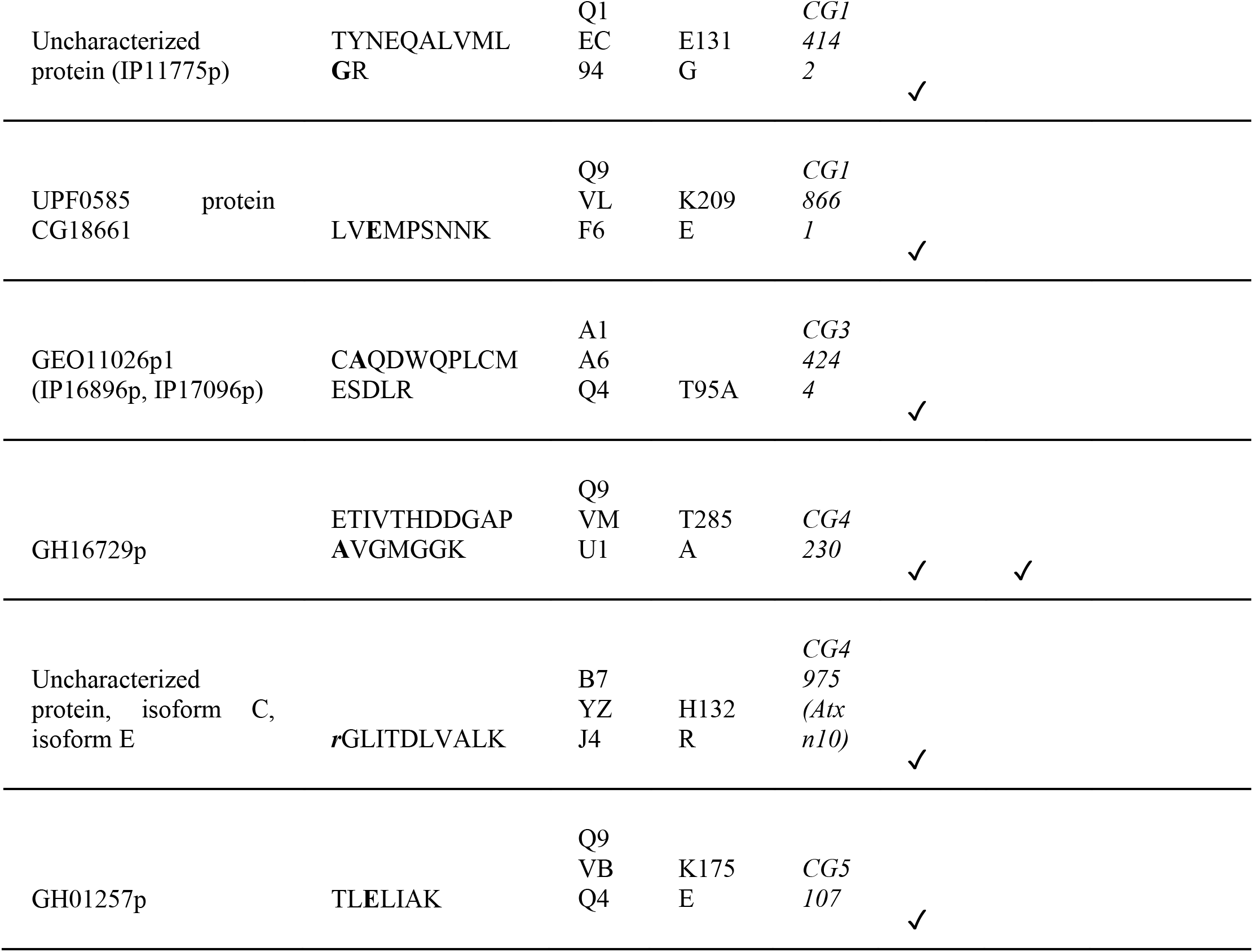
ADAR-edited coding sites identified in fly proteomes [16,31]. The sites were detected in Drosophila datasets representing a proteome at different time points during the life cycle and embryonic development under accession numbers PXD005691 and PXD005713, respectively [31]. The list was filtered as described in Methods. As compared with the fruit fly edited subproteome from our previous work, 14 shared sites were found [16]. For peptides found in cleaved and miscleaved forms, both variants are shown, with asterisks indicating peptides with higher intensity taken for further analysis. Italic bold lowercase letters indicate an editing site located to the right of the peptide and not represented in the peptide itself. More detailed information about identified edited sites is presented in Suppl. Table 1 (life cycle) and Suppl. Table 2 (embryogenesis).

A relatively short list of identified edited proteins did not show a significant pathway enrichment when multiple comparisons were corrected. However, protein-protein interaction analysis using STRING knowledgebase [35] indicated a considerable amount of interactions between the proteins. The knowledgebase returns a p-value of PPI enrichment in the list of edited proteins as low as 0.0004, indicating that proteins have more interactions among themselves than what would be expected by chance. This enrichment means that non-synonymous RNA editing is not stochastic and may modulate work of ensembles of interactors in the need of such a regulation.

Interacting proteins classified into functional groups (Table 1) are represented in Figure 1. One group of functionally connected proteins is involved in actomyosin structure organisation, with titin and paramyosin as its widely known representatives. Further, a group of interacting proteins represents members of vesicle trafficking pathways functionally related to the machinery of synaptic vesicle release. Of those, complexin was added to this group manually, as it was shown earlier to be an important edited component of SNARE complex, which is responsible for the fusion of the presynaptic membrane and synaptic vesicles [16]. Editing of a few interacting SNARE complex components was detected in fruit fly brain proteomes, as described below. The rest of edited proteins, except those which are still not characterized, were classified into smaller groups of cytoskeleton components, participants of translation and proteostasis machinery, proteins involved in RNA processing and binding, egg-specific and chromatin proteins, as well as a single ion channel (Table 1).

**Figure 1.**
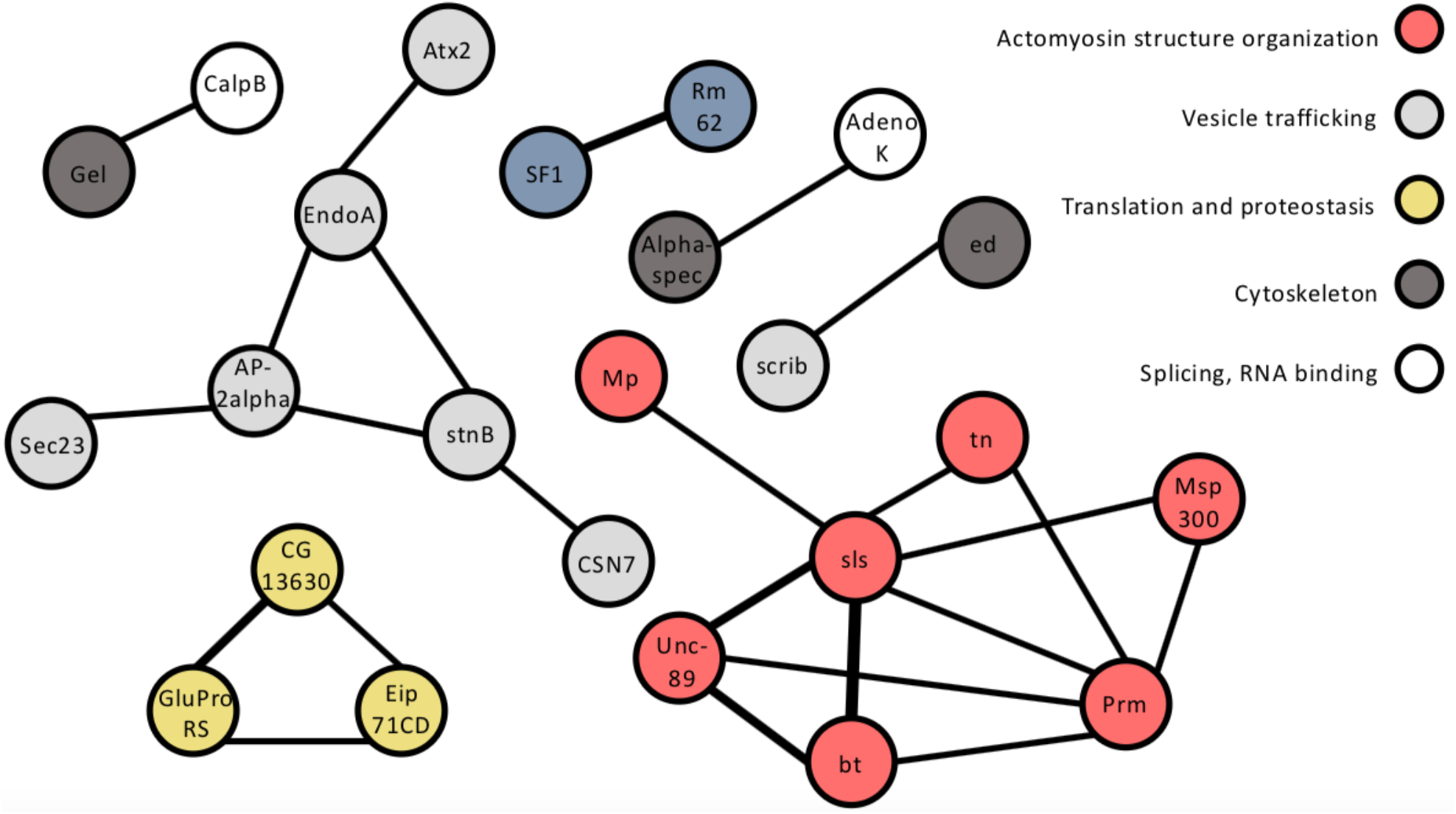
Protein-protein interactions for proteins that were found to undergo RNA editing generated by the STRING knowledgebase [35]. Thickness of lines depicts the number of methods confirming each interaction according to STRING. Colors encodes edited protein subsets as defined in Table 1.

### Distribution of edited proteins along life cycle stages

Using proteomic data, we determined features characterizing the editing levels of found protein edited sites and expression levels of the corresponding proteins, so that these levels between developmental stages could be further compared. All edited sites were clustered using precursor ion (MS1) intensity as a measure of the corresponding tryptic peptide level. Notably, the unsupervised clustering has distributed the edited proteins in remarkably exact agreement with the life cycle phases (Figure 2). One can assume that this distribution reflects the expression levels of respective proteins, as phase-specific proteins were recognized before [31]. However, the same proteins clustered using their expression level estimated using the normalized spectral abundance factor (NSAF) [36] did not show such a good agreement with the life cycle phases (Suppl. Figure 1). Further, these NSAF clusters had different composition compared with MS1 clusters. In most cases, the Pearson correlation coefficient between MS1 intensities of tryptic peptides containing edited sites and NSAF values of the corresponding proteins was not significantly different from zero (p-value greater than 0.05; grey bars on the right, Figure 2). Taken together, these findings clearly indicate that non-synonymous RNA editing modulates sequences of affected proteins and, presumably, their functions in a life cycle phase-specific manner.

**Figure 2.**
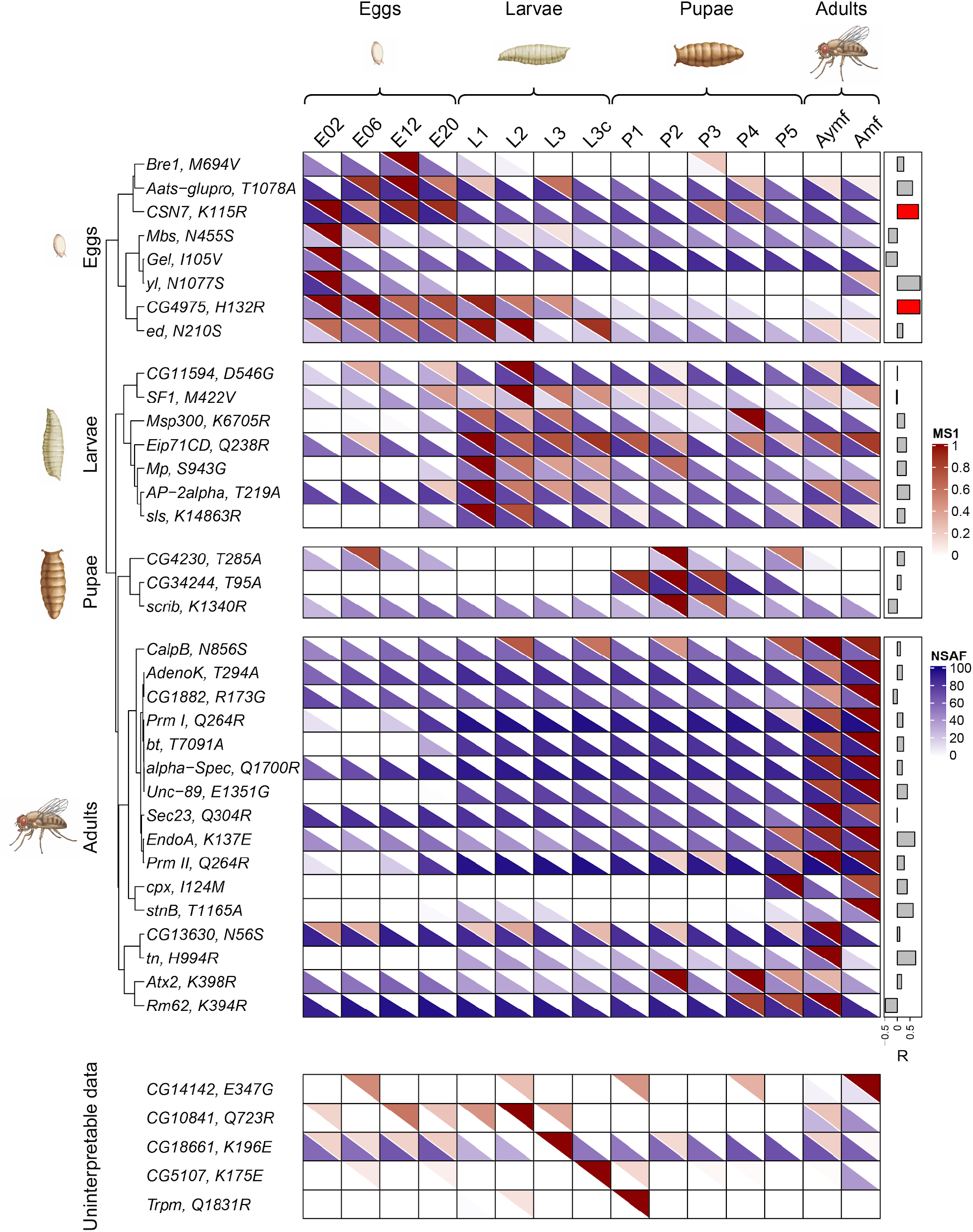
A heatmap of relative abundances of edited sites as tryptic peptides and corresponding proteins at various stages of the fruit fly life cycle. The extent of editing (red half of each cell of the diagram) is expressed in MS1 intensity values for the corresponding variant peptides, normalized for each row individually. The protein abundance (blue half of each cell) is expressed as NSAF percentiles [36]. Pearson correlation coefficients between MS1 and NSAF values are shown on the right. For each coefficient, the p-value of its difference from zero is calculated. Values highlighted in red are Pearson coefficients with FDR adjusted p-value less than 0.05. Clustering was performed using MS1 intensities. MS1 and NSAF values designated by colors on the heatmap are listed in Suppl. Table 1. If the edited site was identified and protein abundance was impossible to estimate, the data were considered uninterpretable. Staging of life cycle was taken from the paper where the primary proteomic data were taken from [31].

We may only hypothesize how this phase-specific editing is achieved, taking into account the single gene producing ADAR proteoforms in the insect. Factors such as modulation of its enzymatic activity by developmental protein regulators and switches of splice isoform [29] deserve more thorough consideration.

A cluster of proteins edited predominantly at the embryonic stage includes an egg-specific vitellogenin receptor encoded by *yl* gene. A subunit of COP9 signalosome complex (*CSN7*) also edited in this stage is shown to be involved both in oogenesis and embryogenesis [37]. Cytoskeleton-associated proteins gelsolin (*Gel*) and echinoid (*ed*) involved in morphogenesis are also of interest in this group. The echinoid protein and an uncharacterized *CG4975* product are edited not only in the egg phase, but also in larvae. Some products edited in the embryonic phase have also a small peak in the phase of adult animals (Figure 2). For this analysis, we studied imagos of both sexes together. When searches were done separately for males and females, it became clear that the egg-specific identifications were related to adult females containing embryos and not present in males (data not shown).

Larval stages are apparently enriched in edited proteins of muscular machinery. They include titin (*sls*), multiplexin (*Mp*) and muscle-specific protein (Msp300). Of three proteins from the pupal cluster, the *scrib* product is remarkable for its participation in the wing morphogenesis in pupae [38].

The biggest cluster of proteins edited in late pupae and imago insects corresponds to the known burst of ADAR editing associated with neural functions in adult animals [29]. The well-known ADAR targets are seen in the cluster, such as proteins related to synaptic vesicle function and turnover [16]. Those are the components of the SNARE complex, complexin (*cpx*) and stoned B (*stnB*), and endophilin A (*EndoA*), a membrane curvature regulator. Further, an RNA-binding *Atx2* product may represent a link between RNA editing and regulation of circadian rhythms [39]. In addition to neural components, the edited muscular proteins, such as abba/thin (*tn*), paramyosin (*Prm*) and projectin (*bt*), may reflect involvement of RNA editing in formation of adult musculature.

Another dataset from the benchmark paper [31] which describes embryogenesis of fruit fly in detail yielded much less edited sites than a major life cycle dataset, as shown above in Table 1. Clustering twelve sites in the same manner as in Figure 2 expectedly failed to define any patterns (see Suppl. Figures 2-3) due to the low amount of available components. In these data, levels of three edited sites correlated with the protein abundances.

### ADAR-edited sites in transcriptomic and proteomic data for the fruit fly life cycle

It was interesting to compare proteomic consequences of A-to-I RNA editing with the transcriptomic data available for *D. melanogaster* life cycle. In one of the most comprehensive studies involving 30 time points in insect lifespan, 972 ADAR editing events were monitored transcriptome-wide [22]. According to the previous works, the transcriptomic data demonstrated that the editing in most sites started at the late pupal stage, despite some transcripts being edited in early embryo [22]. Of the transcriptomic results, we considered 624 non-synonymous events to compare them with 42 edited sites obtained from the reanalyzed proteomic data above. We found only 12 sites being common for two datasets at the end (Suppl. Table 3). This low coincidence was not surprising and may be explained by both biological and technical factors. Indeed, it was shown previously that the correlation between abundances of mRNA transcripts and corresponding protein products may be low due to different lifespans of these molecules in cells and body fluids [40]. The techniques of transcriptomics and proteomics which differ in sensitivity, among the other analytical features, also cause the disparity in the results. A multi-omic approach where the same specimens are used to isolate RNA and proteins is expected to improve reproducibility, although it still has not been employed in the RNA editing studies.

A heatmap which compares editing percentages of transcripts with relative abundances of the corresponding sites in proteins is shown in Figure 3. The first notable thing is that only the late edited sites, which originate from RNA editing burst in the late pupal or adult phases, have their transcriptomic counterparts. Editing of transcripts, in most cases, was detected a few stages earlier than the same for proteins. This may be explained by the limited sensitivity of mass spectrometry to detect peptides of interest in comparison with amplification of nucleic acids which can facilitate detection of a single molecule of transcript. Gradually, with the increase in abundance of edited proteoform, it will become detectable at the protein level.

**Figure 3.**
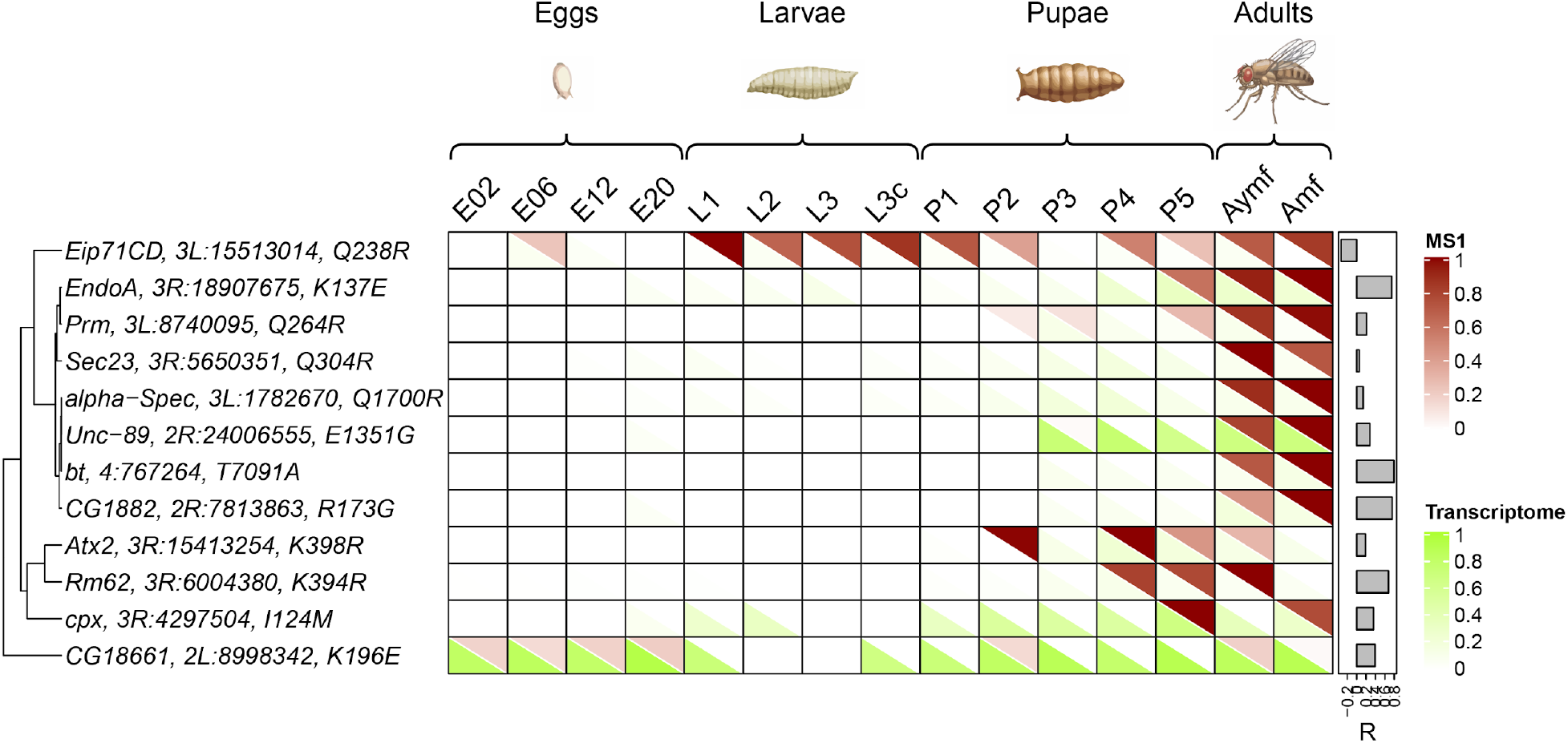
Abundances of edited protein sites and the corresponding sites from edited transcripts at various stages of fruit fly life cycle. The protein editing extent (red upper half of each rectangle) is expressed as MS1 intensities of the corresponding peptides normalized for each row individually. The transcript abundance (green lower half of each rectangle) is expressed as the editing percentage taken from the earlier work [22]. Data are clustered based on Pearson correlation coefficients between MS1 and editing percentage values shown on the right. For each coefficient, the p-value of its difference from zero is calculated which was higher than 0.05 in all cases (no significant correlation). Clustering was performed using MS1 intensities. MS1 and editing percentage values designated by colors on the heatmap are listed in Suppl. Table 1b and Suppl. Table 3, respectively. A single outlier value of MS1 was excluded from the data for CG18661 gene. Staging of life cycle was taken from [31].

### Quantitation of brain-specific edited proteins during a fruit fly life cycle

Following the analysis of edited protein sites from whole bodies, we performed studies on fruit fly brain editome. Inventory of edited protein sites characteristic for this organ was done before for adult insects [16]. In that work, the targeted proteomic assays to measure edited and genomically encoded sites in selected proteins were suggested based on their repeated identification as edited in brain and head proteomes, as well as their relation to neuronal functions [16]. Then, we quantified the editing levels of these protein sites in brains isolated from *D. melanogaster* at different life cycle phases, from larvae through pupae to imagos.

The edited proteins of interest are mostly related to synaptic signaling and, particularly, to pathways that mediate a fusion of a synaptic vesicle with a presynaptic membrane for release of the former [41]. Among them, syntaxin 1A (*Syx1A*) and complexin (*cpx*) represent a SNARE presynaptic complex responsible for vesicle-membrane fusion [42]. Endophilin A (*EndoA*) regulates membrane curvature, including this parameter for synaptic vesicles [43]. *Cadps* and *CG4587* gene products participate in calcium-dependent signaling in the presynaptic zone [44,45]. In addition, an edited site of ataxin-2 homolog (Atx2) was included in the list. This protein is a homolog of a human polyglutamic disease factor [46] and was shown to be involved in various processes, such as eye development, actin filament formation and circadian activity regulation [47,48].

Edited and genomically encoded sites of six selected proteins were quantified in brains of *D. melanogaster* at three life cycle phases using targeted mass spectrometry based on multiple/selected reaction monitoring with stable isotope standards (MRM-SIS). The standards were designed and tested previously to measure protein editing in adult insect brains [16]. Herein, we repeated all biological experiments in two independent fly cultures and, for technical reasons, used different chromatography-mass spectrometry methods, which are described in more detail in the Methods section. These experiments allowed estimation of the interference between technical variability and quantitation results. The data from all biological and technical replicates are listed in Suppl. Table 4.

The MRM-SIS approach is intended to provide absolute quantitation of proteins of interest. However, the method measures the proteolytic peptide concentrations, and technical variance may be introduced during protein isolation from the sample and digestion by protease. A real absolute quantitation by the method is reached when all experimental steps are highly standardized including robotisation and solving the metrological issues [49]. In our case, there was a difference between absolute quantitation data normalized by the total protein content obtained for different biological replicates (Suppl. Figure 4). At the same time, the results were consistent within each replicate and relative levels of measured products were reasonably convergent. We expressed the rate of each site editing as a percentage of edited product (Figure 4), in accordance with a form of representation established for RNA sequencing results to estimate the same on the RNA level [50].

Out of six edited protein sites considered in this study, five sites yielded quite interpretable results (Figure 4). Unfortunately, both edited and genomic forms of the Atx2 homolog were found in a single biological replicate near the limit of detection, and it was difficult to characterize them during the life cycle. The rest of the sites demonstrated a trend towards increasing the editing rate from larval to adult brain (Figure 4). A Lys-137-Glu site of endophilin A was the most edited one reaching 75-80% of editing in the adult brain. At the same time, the editing rate for this site was also relatively high even in larvae. As we previously hypothesized, the substitution may recharge a membrane-contacting surface of endophilin A dimer and decrease the affinity of its binding to the intracellular membranes due to change in electrostatic interactions [16].

**Figure 4.**
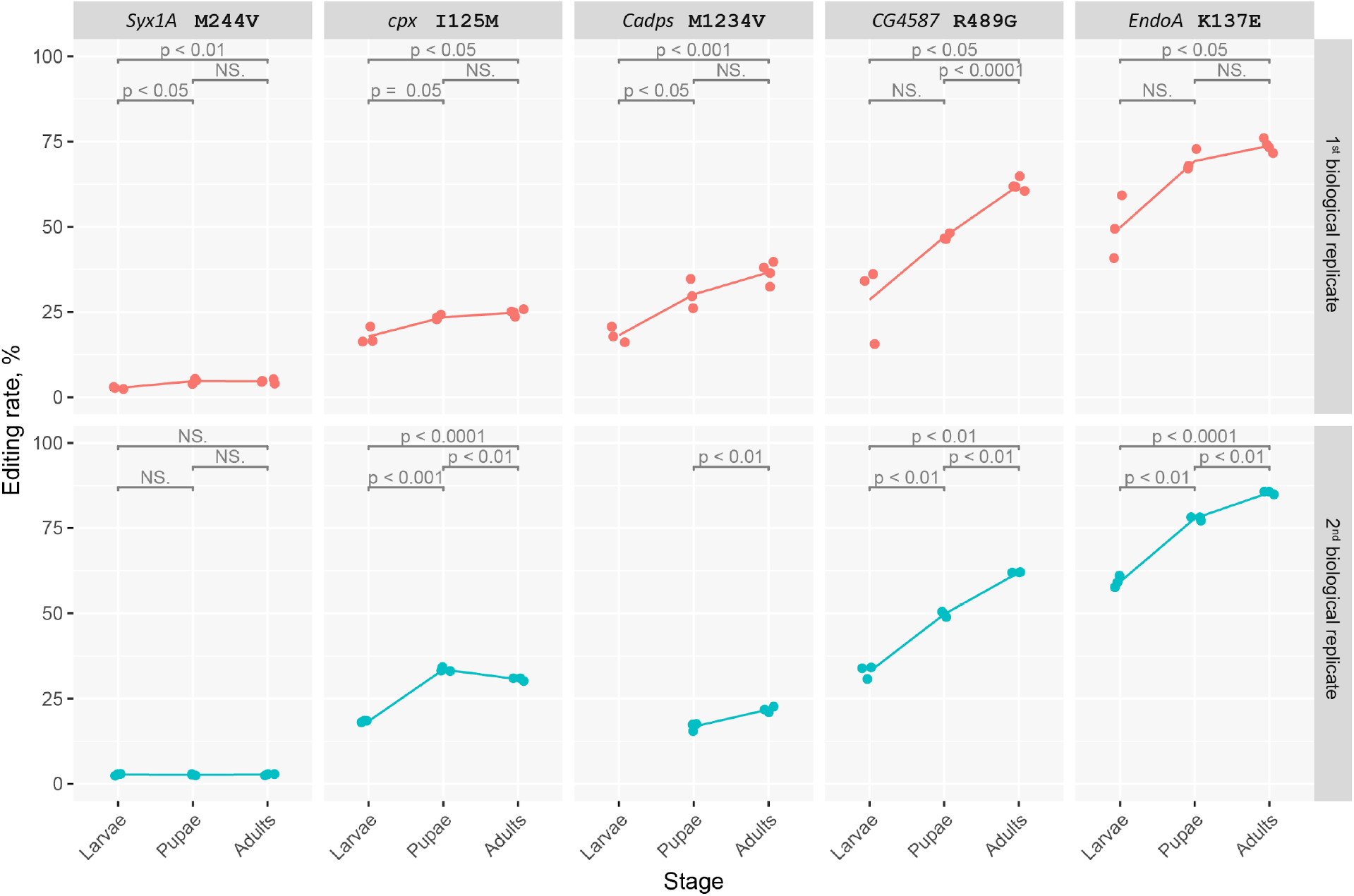
Dynamics of editing rates for selected protein sites in the brain at life cycle phases of *D. melanogaster*. Data is shown independently for two biological replicates. The editing rate was calculated as the concentration of the edited variant divided by the sum of concentrations of both variants. T-test’s p-values are presented to estimate the difference in the editing level between life cycle phases. For *Cadps*, in the second replicate, the edited site was not detected at the larval phase.

Of two components of SNARE complex [42], editing of complexin (*cpx*) site significantly increased from larval to pupal stages and then stabilized or even slightly lowered. As for syntaxin 1A (*Syx1A*), its editing rate was low at all stages with a trend to increase in one of biological replicates.

Calcium-dependent secretion activator is a protein that is functionally close to the SNARE complex, as it also promotes calcium-dependent vesicle release [44]. Its moderately edited site is also characterized by increasing editing rate during life cycle. Notably, this protein is one of the few, which are also edited by ADAR in mammals. However, its edited sites found in human and murine proteomes are not conservative and differ from Met-1234-Val from fruit fly [16]. Another protein, which is connected with calcium signaling for neurotransmission, is the voltage-gated calcium channel encoded by *CG4587*. Non-synonymous RNA editing of its orthologs which is able to modulate calcium influx through the channels is widely recognised in rodents [51], although the Arg-489-Gly substitution reported here for fruit fly is not conserved as an edited site in mammals. Editing of this site is extensively growing at each phase of life cycle, presumably illustrating its functional importance. This type of substitution from RNA editing changes the size and electrostatic charge of a residue and was shown to modulate their electrophysiological properties for rat glutamate channel subunits [52].

In agreement with the transcriptomic data, targeted analysis of neural edited proteins illustrated that insect metamorphosis is accompanied by an increase in editing rate, an editing burst described earlier [29] and observed in all data above. Notably, at least five of six proteins were edited at the larval phase, further supporting the existence of a basal level of ADAR activity in early development.

Soon after its discovery [53], ADAR-mediated RNA editing in model organisms was studied using inhibiting or modifying ADAR activity by genetic mutations and observing effects on different levels of body organization. The fruit fly is one of the oldest and well established model species, which provides a suitable source of mutants for functional characterization of RNA editing. Recently, some mechanisms resulting in neurodegeneration in the absence of ADAR activity were discovered using *D. melanogaster* strain with inactivated enzyme [28]. Further, a point mutation was induced in the fruit fly ADAR gene, which preserved an expression level of the resultant protein, but inhibited its enzymatic activity [13]. Using these model animals, a unique binary function of Drosophila’s ADAR was shown, which shared features of mammalian ADAR1 participating in innate immunity suppression and, at the same time, ADAR2 editing mRNA and, consequently, protein sequences [13].

In addition to the functionality studies using mutated models, the molecular effects of ADAR activity in the fruit fly were explored transcriptome-wide [54,55] and, more recently, proteome-wide [16]. As mentioned above, a fatal ADAR2 deficit in mice can be reversed by genomic editing of a single site in GRIA2 glutamate receptor subunit [19]. However, these results cannot be easily extended on the human health issues because of different lifespan and physiology in rodents. Further, a recent comprehensive study of fruit fly mutants illustrated that, at least, the locomotory deficit associated with impairment of ADAR enzymatic function is mediated by edited proteins [13]. At the same time, it remains unknown how exactly edited proteins modulate cellular and physiological functions. Limited works on mice demonstrated an approach to study functional significance of selected coding edited events by point knock-out or knock-in sites of interest in the model genome, such as *Gria2* [19] or *Flna* [20] mutants.

Proteome-wide identification of proteoforms caused by RNA editing provides an instrument for elucidation of the roles of certain edited sites in the wild-type and mutant *Drosophila* strains. We provided such data for developmental phases of fruit fly lifespan. The results demonstrated that phase-specific editing of proteins may exist independently of their expression levels. However, it remains unclear which factors regulate this editing during the life cycle. In accordance with the prior transcriptomic data, we also observed at the proteome level the burst in nonsynonymous editing by ADAR during insect metamorphosis. Positive dynamics of protein editing was validated with targeted mass-spectrometry in fruit fly brain proteins which was one of the first examples when protein editing extent was estimated quantitatively.

## Materials and methods

### RNA editome database for proteomics search

The database was formed as described before [16]. Briefly, fly’s genomic coordinates of RNA editing sites mapped to exons were obtained from RADAR database (1328 sites) [56] and genome-wide study by St Laurent et al [54]. The resultant database contained about eight thousands of non-synonymous editing sites. Genomic coordinates obtained from these sources were converted to the coordinates in the Drosophila genome assembly Dm6 using FlyBase (http://flybase.org/) [57]. Changes in protein sequences induced by RNA editing were annotated using Variant Annotation Integrator (VAI; http://genome.ucsc.edu/cgi-bin/hgVai).

### Proteomic data for fruit fly ontogeny selected for reanalysis

Proteome datasets were taken from ProteomeXchange repository [58]. The datasets represent shotgun proteomes for embryonic development (PXD005713) and the whole life cycle of fruit fly (PXD005691) [31]. In these datasets, proteomes of the whole bodies of *D. melanogaster* Oregon R line grown at 25°C were studied. The embryogenesis dataset was obtained from eggs collected every hour in the range from 0 to 6 hours and then every 2 hours up to 20 hours. The whole life cycle was analyzed in the following stages: zygotic gene activation (0-2 hours), gastrulation (4-6 hours), organogenesis (10-12 hours) and the late stages of embryogenesis (18-20 hours). For larva, four stages were chosen: L1, L2, early L3, late L3 (crawling larva). Pupae were analyzed at the following time points: P1 (0-14 hours after pupation), P2-P5 for every 24 hours after pupation. Of adult data, the virgin flies (adult young male and female - *aymf*) and one week old animals (adult male and female - *amf*) were selected for analysis.

### Proteomic search and output filtering

RAW files were converted to mzML using MSConvert [59]. All files were then searched with IdentiPy [60] using the following parameters: fixed carbamidomethylation of cysteine and variable oxidation of methionine; auto-tuning of parameters enabled, with 10 ppm and 0.01 Da as initial values for precursor and fragment ion mass tolerances; Dinosaur [61] was called from IdentiPy automatically to deconvolute chimeric spectra and extract MS1-level intensity information. Search results were post-processed with Scavager [62] in two modes: first pass to obtain the lists of validated PSMs, peptides and proteins filtered to 1% FDR and a second one for group-specific filtering of identifications containing RNA editing sites. For group-specific FDR filtering, FDR correction [63] was disabled, allowing for higher true values of FDR. Identifications from all run files corresponding to each development stage (see Suppl. Tables 1-2) were pooled at PSM level for postprocessing.

After that, a table of peptides containing RNA editing sites and identified in at least one sample was constructed for each of the datasets. As a quantitative measure for identified peptides we used MS1 intensity extracted with Dinosaur. To quantify the wild-type proteins corresponding to these peptides we used NSAF values as reported by Scavager and converted to percentage ranks to eliminate systematic shifts between samples. The peptide table contained the following information on each peptide: genomic coordinates of its RNA editing site, PSM count and max MS1 intensity in each sample.

Primary identifications of edited sites were filtered to avoid reporting false positive hits as described earlier [16]. First, the results were manually checked for possible editome database errors. Then, tryptic peptides containing Asn-to-Asp (N>D) substitution were excluded because this change is not distinguishable from natural deamidation of asparagine. Based on our previous finding [16], solitary miscleaved peptide identifications reported without their properly cleaved counterparts are enriched in false positives. Thus, hits of this type were also excluded from results. Further, the identifications supported by a single PSM in all data were considered as dubious and not reported in the main text. Those latter can be found in Suppl. Tables 1-2.

Additionally, DeepRT algorithm was applied to all peptides [64]; predicted retention times were linearly transformed to align with experimental times in each experimental run separately, using top 200 PSMs (by score) from every file to find the optimal linear transformation. Difference between predicted and experimental retention times (RT) was converted to Z-score for each PSM by normalizing it with the standard error of the linear regression fitting RTs predicted with DeepRT to experimental ones. Hits with Z-score > 2.5 were then filtered out. High values of Z-score corresponded to outliers on the experimental and predicted RT plotted as a linear correlation (Suppl. Table 5 and Suppl. Figure 5 exemplified for the dataset PXD005691).

### Protein interactions and data visualization

Protein-protein interactions were obtained from STRING resource [35]. Minimum required interaction score was set to medium confidence (0.4), whole genome used as a statistical background.

Heatmap rows were arranged according to hierarchical complete linkage clustering. Pearson correlation between MS1 intensity values was utilized as distance metric (defined as 1−R). Dendrogram rows were reordered with weights proportional to MS1 value and development stage when the protein was detected. Heatmaps were plotted using the ComplexHeatmap package [65].

### Animals

A culture of *D. melanogaster* Canton S line was used for targeted proteomics analysis. The flies were kindly provided by Dr. Natalia Romanova (Moscow State University, Department of Biology). The line was kept on Formula 5-24 instant Drosophila medium (Carolina Biological Supply Company, USA) in 50 mL disposable plastic test tubes.

For targeted proteomic analysis, brains of third stage larvae (L3), late pupae (P12-14 according to [66]) and young adult flies (approximately 7 days old) of both sexes were used. The culture was maintained in the dark at 25°C. Brain dissection for all phases was performed as described previously [16]. Briefly, alive insects were taken out, and the body was rapidly removed by a needle. The head was placed into 0.01 M PBS at pH 7.4 (Sigma-Aldrich, USA), and the head capsule was torn apart by two forceps under visual control through a stereo microscope (Nikon SMZ645, Japan) with 10 × 1 magnification. The extracted brains were collected into the same buffer solution, and then centrifuged at 6000 g for 15 min (Centrifuge 5415R, Eppendorf, Germany). PBS was removed, and the brain pellet was frozen at −80 °C until sample preparation. Two biological replicates were obtained independently with a time gap of six months.

### Brain sample preparation

For the first biological replicate, 80 brains were used to obtain each sample. For each stage of the second replicate, 100 brains were lysed. The rest of the sample preparation protocol was similar for both replicates.

Frozen fly brains were resuspended in 100 μL of lysis solution containing 0.1% (w/v) ProteaseMAX Surfactant (Promega, USA), 50 mM ammonium bicarbonate, and 10% (v/v) acetonitrile (ACN). The cell lysate was stirred for 60 min at 550 rpm at room temperature (TS-100, BioSan, Latvia). Then the mixture was subjected to sonication by Bandelin Sonopuls HD2070 ultrasonic homogenizer (Bandelin Electronic, Germany) at 30% amplitude using short pulses for 5 min. The supernatant was collected after centrifugation at 15,700g for 10 min at 20 °C (Centrifuge 5415R, Eppendorf, Germany). Total protein concentration was measured using bicinchoninic acid assay (BCA Kit, Sigma-Aldrich, USA).

A total of 2 μL of 500 mM dithiothreitol (DTT) in 50 mM triethylammonium bicarbonate (TEABC) buffer was added to the samples to the final DTT concentration of 10 mM followed by incubation for 20 min at 300 rpm at 56 °C. Thereafter, 2 μL of 500 mM iodoacetamide (IAM) in 50 mM TEABC were added to the sample to the final IAM concentration of 10 mM. The mixture was incubated in the darkness at room temperature for 30 min.

The total resultant protein content was digested with trypsin (Trypsin Gold, Promega, USA). The enzyme was added at the ratio of 1:40 (w/w) to the total protein content, and the mixture was incubated overnight at 37 °C. Enzymatic digestion was terminated by addition of acetic acid (5% w/v). Then, the samples were stirred at 500 rpm for 30 min at 45 °C followed by centrifugation at 15,700g for 10 min at 20 °C. The supernatant was then added to the filter unit (30 kDa, Millipore, USA) and centrifuged at 13,400g for 20 min at 20 °C. After that, 100 μL of 50% formic acid was added to the filter unit and the sample was centrifuged at 13,400g for 20 min at 20 °C. The final peptide concentration was measured using Peptide Assay (Thermo Fisher Scientific, USA) on a NanoDrop spectrophotometer (Thermo Fisher Scientific, USA). The sample was dried using a vacuum concentrator (Concentrator 5301, Eppendorf, Germany) at 30 °C. Dried peptides were stored at −80 °C until the LC-MS/MS analysis.

### Synthesis of stable isotope-labeled peptide standard

Peptides were synthesized by solid phase method using amino acid derivatives with 9-fluorenyl methyloxy carbonyl (Fmoc) protected α-amino groups (Novabiochem). The procedure was performed as described previously [16]. Stable isotope containing leucine (Fmoc-Leu-OH-13C_6_,^15^N, Cambridge Isotope Laboratories) was applied for labeling 11-plex peptides from *cpx* protein (NQMETQVNE***L*** ^h^K and NQIETQVNE***L*** ^h^K). Resin with attached stable isotope-labeled lysine (L-Lys (Boc) (^13^C_6_, 99%; ^15^N_2_, 99%) 2-Cl-Trt, Cambridge Isotope Laboratories) was used for synthesis of peptides of *Syx1A* (IEYHVEHAMDYVQTATQDT***K*** ^h^ and IEYHVEHAVDYVQTATQDT***K*** ^h^), *EndoA* (YSLDDNI***K*** ^h^ and YSLDDNIEQNFLEPLHHMQT***K*** ^h^), *Cadps* proteins (LMSVLESTLS***K*** ^h^ and LVSVLESTLS***K*** ^h^), and one of the peptides of *Atx2* (GVGPAPSANASADSSS***K*** ^h^). Resin with attached stable isotope-labeled arginine (L-Arg (Pbf) (^13^C_6_, 99%; ^15^N_4_, 99%) 2-Cl-Trt, Cambridge Isotope Laboratories) was used for synthesis of peptides of *CG4587* (LVTTVSTPVFD***R*** ^h^ and LVTTVSTPVFDG***R*** ^h^) and *Atx2* (GVGPAPSANASADSSS***R*** ^h^). Further steps of synthesis were also performed as described earlier [67].

For quality control of the synthesis, LC-MS analysis was done using a chromatographic Agilent ChemStation 1200 series connected to an Agilent 1100 series LC/MSD Trap XCT Ultra mass spectrometer (Agilent, USA). Since some peptides contained methionines, the quality control also included manual inspection of the MS and MS/MS spectra for possible presence of the peaks produced by oxidized compounds. No such peaks were found in our case.

Absolute concentrations of synthesized peptides were determined using conventional amino acid analysis with their orthophthalic derivatives according to standard amino acid samples.

### Quantitative analysis by multiple reaction monitoring using stable isotope standards

Each sample for the first biological replicate was analyzed using Dionex UltiMate 3000 RSLC nano System Series (Thermo Fisher Scientific, USA) connected to a triple quadrupole mass spectrometer TSQ Vantage (Thermo Fisher Scientific, USA) in three technical replicates for larval and pupal stages and four replicates for imago. Generally, 1 μl of each sample containing 2 μg of total native peptides and 100 fmol of each standard peptide was loaded on a precolumn Zorbax 300SB-C18 (5 μm, 5 × 0.3 mm) (Agilent Technologies, USA) and washed with 5 % acetonitrile for 5 min at a flow rate of 10 μl/minute before analytical LC separation. Peptide separation was carried out on an RP-HPLC column Zorbax 300SB-C18 (3.5 μm, 150 mm × 75 μm) (Agilent Technologies, USA) using a linear gradient from 95 % solvent A (0.1 % formic acid) and 5 % solvent B (80 % acetonitrile, 0.1 % formic acid) to 60 % solvent A and 40% solvent B over 38 minutes at a flow rate of 0.3 μl/minute. Multiple reaction monitoring (MRM) analysis was performed on QqQ TSQ Vantage (Thermo Scientific, USA) equipped with a nano-electrospray ion source. The set of transitions is reported in Suppl. Table 6a. Capillary voltage was at 2100 V, isolation window was set to 0.7 Da for the first and the third quadrupole, respectively, cycle time was 3 s. Fragmentation of precursor ions was performed at 1.0 mTorr, using collision energies calculated with Skyline v. 3.1 [68] software.

Measurements for the second biological replicate were performed as follows. Each sample was analyzed using a microflow Agilent 1200 HPLC system (Agilent Technologies, USA) connected to a triple quadrupole mass spectrometer TSQ Quantiva (Thermo Fisher Scientific, USA) in three technical replicates. Generally, 3 μl of each sample containing approximately 10 μg of total native peptides and 500 fmol of each stable isotope-labeled standard (SIS) peptide were loaded on an analytical column, Zorbax 300SB-C18 (5 μm, 150 × 0.3 mm) (Agilent Technologies, USA) and washed with 5% acetonitrile for 5 min at a flow rate of 20 μl/min. Peptides were separated using a linear gradient from 95% solvent A (0.1% formic acid) and 5 % solvent B (80% acetonitrile, 0.1% formic acid) to 50% solvent A and 50% solvent B over 30 minutes at a flow rate of 20 μl/minute. MRM analysis was performed using triple quadrupole TSQ Quantiva (Thermo Scientific, USA) equipped with an electrospray ion source. The set of transitions used for the analysis is shown in Suppl. Table 6b. Capillary voltage was set at 4000 V, ion transfer tube temperature was 325 °C, vaporizer temperature was 40 °C, sheath and aux gas flow rates were set at 7 and 5 L/min, respectively. Isolation window was set to 0.7 Da for the first and the third quadrupole, and the cycle time was 1.2 s. MRM experiment was performed in a time-scheduled manner with retention time window of 2 min for each precursor ion. Fragmentation of precursor ions was performed at 1.2 mTorr using collision energies calculated by Skyline 4.1 software (MacCoss Lab Software, USA) [68]. The need for recalculation of parameters for analysis was dictated by use of another mass-spectrometer for the second biological replicate.

Quantitative analysis of MRM data was performed using Skyline 4.1 software. Quantification data obtained from the "total ratio" numbers calculated by Skyline represented a weighted mean of the transition ratios, where the weight was the area of the internal standard. Up to five transitions were used for each peptide including the isotopically labeled standard peptide. Isotopically labeled peptide counterparts were added in amounts of 50 fmol/mkg of total protein. Each MRM experiment was repeated in 3 technical runs. The results were inspected using Skyline software to compare chromatographic profiles of endogenous peptide and stable-isotope labeled peptide. CV of transition intensity did not exceed 20% in technical replicates.

## Supporting information

Suppl. Table 1. A-to-I-edited peptide hits reported by IdentiPy from reanalyzed data of ProteomeXchange dataset PXD005691.

Suppl. Table 2. A-to-I-edited peptide hits reported by IdentiPy from reanalyzed data of ProteomeXchange dataset PXD005713.

Suppl. Table 3. ADAR-edited sites in transcriptomic and proteomic data.

Suppl. Table 4. Quantitation of selected edited sites and their genomic variants in the brain hydrolysate of the fruit fly by MRM.

Suppl. Table 5. Predicted and experimental retention times (RT) for each PSM obtained from proteome datasets (PXD005691, PXD005713)

Suppl. Table 6. Transition list for MRM method.

## Data availability

All MRM data are deposited to PASSEL (http://www.peptideatlas.org/passel/) [69] under the accession number PASS01553.

## Authors’ contributions

**Anna A. Kliuchnikova:** Visualization, Investigation, Data Curation. **Anton O. Goncharov:** Visualization, Investigation, Data Curation. **Lev I. Levitsky:** Software, Data Curation, Writing - Reviewing and Editing. **Mikhail A. Pyatnitskiy:** Software, Formal analysis. **Svetlana E. Novikova:** Methodology, Investigation. **Ksenia G. Kuznetsova:** Investigation, Data Curation. **Mark V. Ivanov:** Software. **Irina Y. Ilina:** Methodology, Investigation. **Tatyana E. Farafonova:** Investigation, Resources. **Victor G. Zgoda:** Methodology, Resources. **Mikhail V. Gorshkov:** Data Curation, Writing - Reviewing and Editing. **Sergei A. Moshkovskii:** Conceptualization, Methodology, Writing - Original Draft.

## Acknowledgements

The work was supported by Russian Foundation for Basic Research, grant #18-29-13015 to S.M. We thank the Center for Precision Genome Editing and Genetic Technologies for Biomedicine, Federal Research and Clinical Center of Physical-Chemical Medicine of the Federal Medical Biological Agency for providing computational resources for this project. Mass spectrometric measurements were performed using the equipment of the Human Proteome Core Facility of the Orekhovich Institute of Biomedical Chemistry (Russia).

