## Supplemental Figures for "Proteome-Wide Analysis of ADAR-mediated Messenger RNA Editing During Fruit Fly Ontogeny"

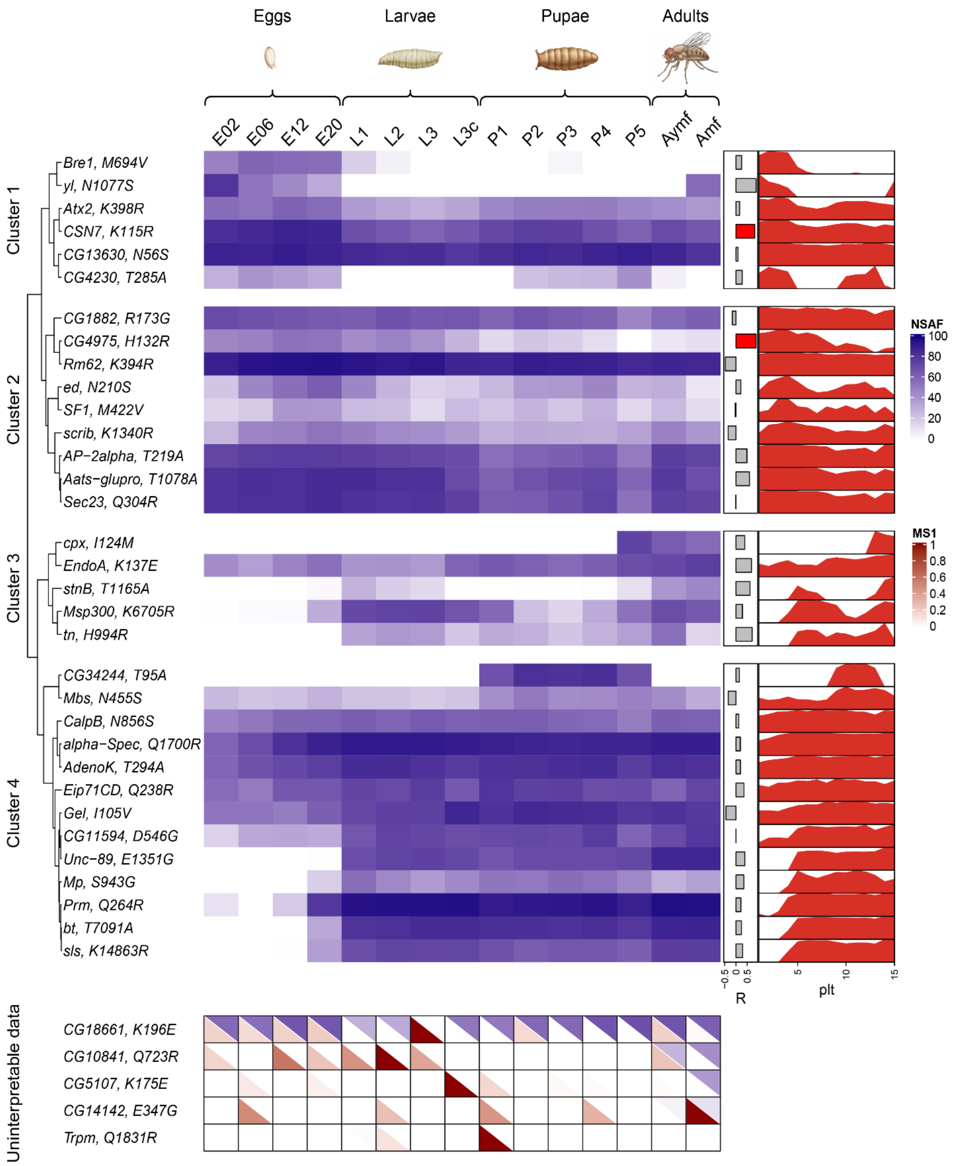


**Supplementary Figure 1. A heatmap representing data for abundance of proteins with editing events at various stages of fruit fly life cycle.** The protein abundance shown in blue is expressed as NSAF percentiles [36]. The Pearson correlation coefficients (R) between NSAF and MS1 (abundance of corresponding edited sites) are represented on the right side as gray and red rectangles. For each coefficient, the p-value of its difference from zero is calculated. The rectangles highlighted in red represents R with FDR adjusted p-value less than 0.05. Clustering was performed by NSAF values. Red plots on the right also represent the change in the edited protein abundance along stages of the life cycle.

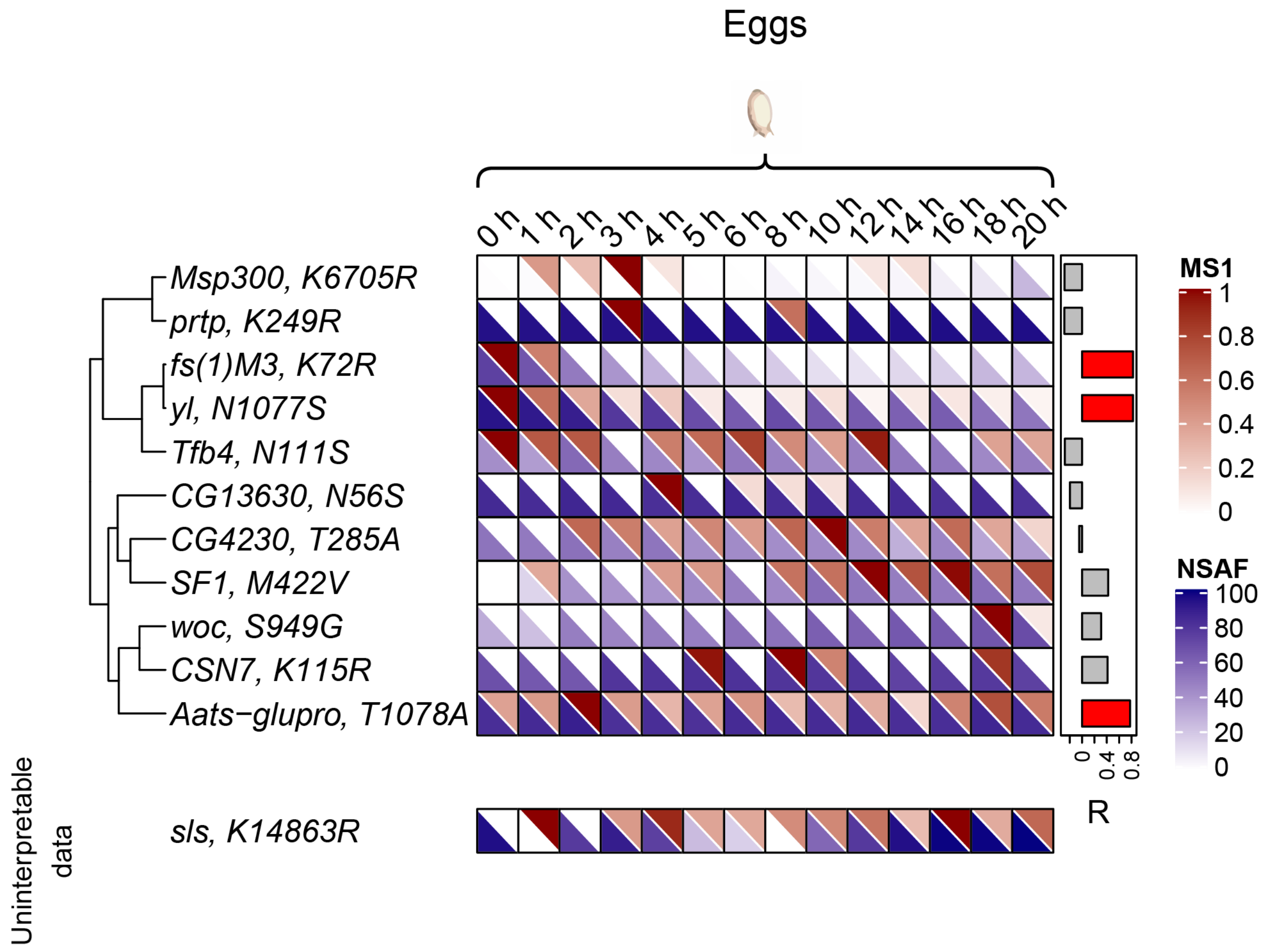


**Supplementary Figure 2**. **A heatmap of relative abundances of edited sites as tryptic peptides and corresponding proteins at various stages of the fruit fly embryogenesis.** The extent of editing (a red half of each cell of the diagram) is expressed in MS1 intensity values, normalized for each row individually. The protein abundance (a blue half of each cell) is expressed as NSAF percentiles [36]. Pearson correlation coefficients between MS1 and NSAF values are represented on the right side. For each coefficient, the p-value of its difference from zero is calculated. Values highlighted in red are Pearson coefficients with FDR adjusted p-value less than 0.05. Clustering was performed by MS1 intensity values. MS1 and NSAF values designated by colors on the heatmap are listed in Suppl. Table 2. If the edited site was identified and protein abundance was impossible to estimate, the data were considered uninterpretable

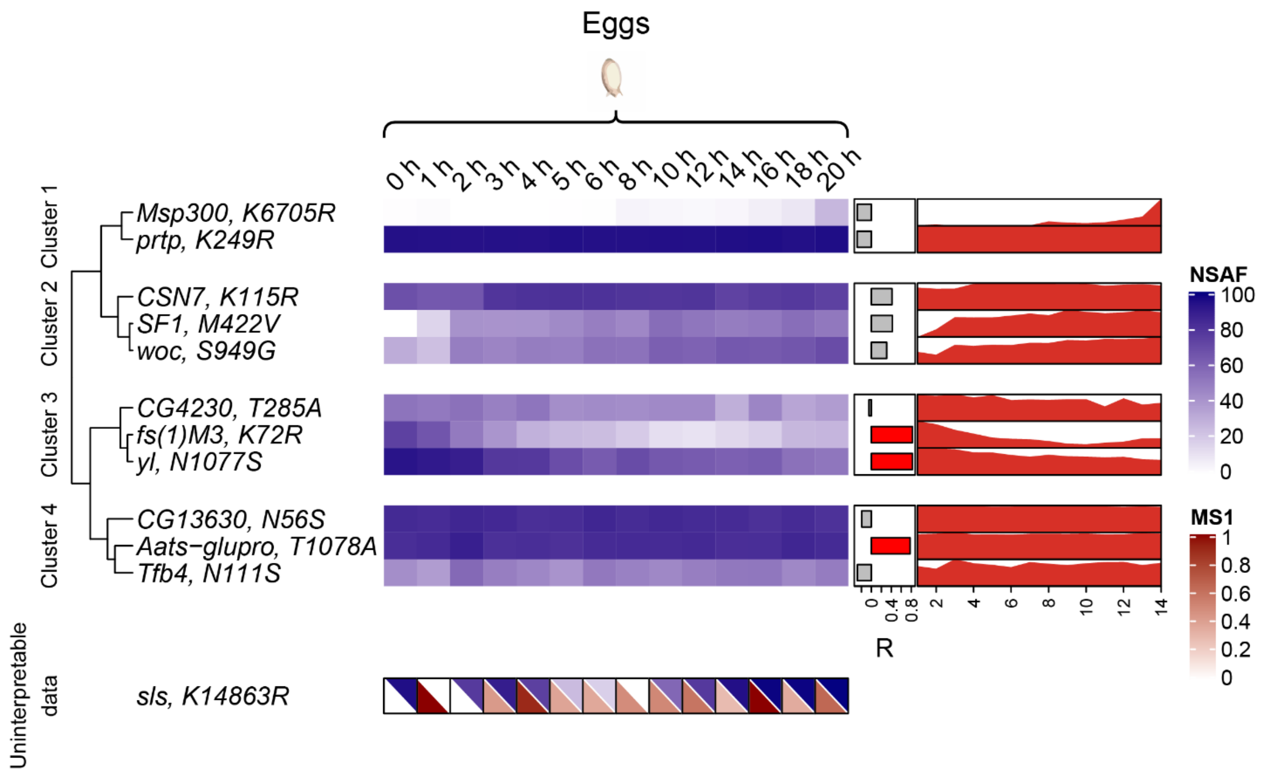


**Supplementary Figure 3**. **A heatmap representing data for abundance of proteins with editing events at various stages of fruit fly embryogenesis.** The protein abundance shown in blue is expressed as NSAF percentiles [36]. The Pearson correlation coefficients (R) between NSAF and MS1 (abundance of corresponding edited sites) are represented on the right side as gray and red rectangles. For each coefficient, the p-value of its difference from zero is calculated. The rectangles highlighted in red represents R with FDR adjusted p-value less than 0.05. Clustering was performed by NSAF values. Red plots on the right also represent the change in the edited protein abundance along stages of the life cycle.

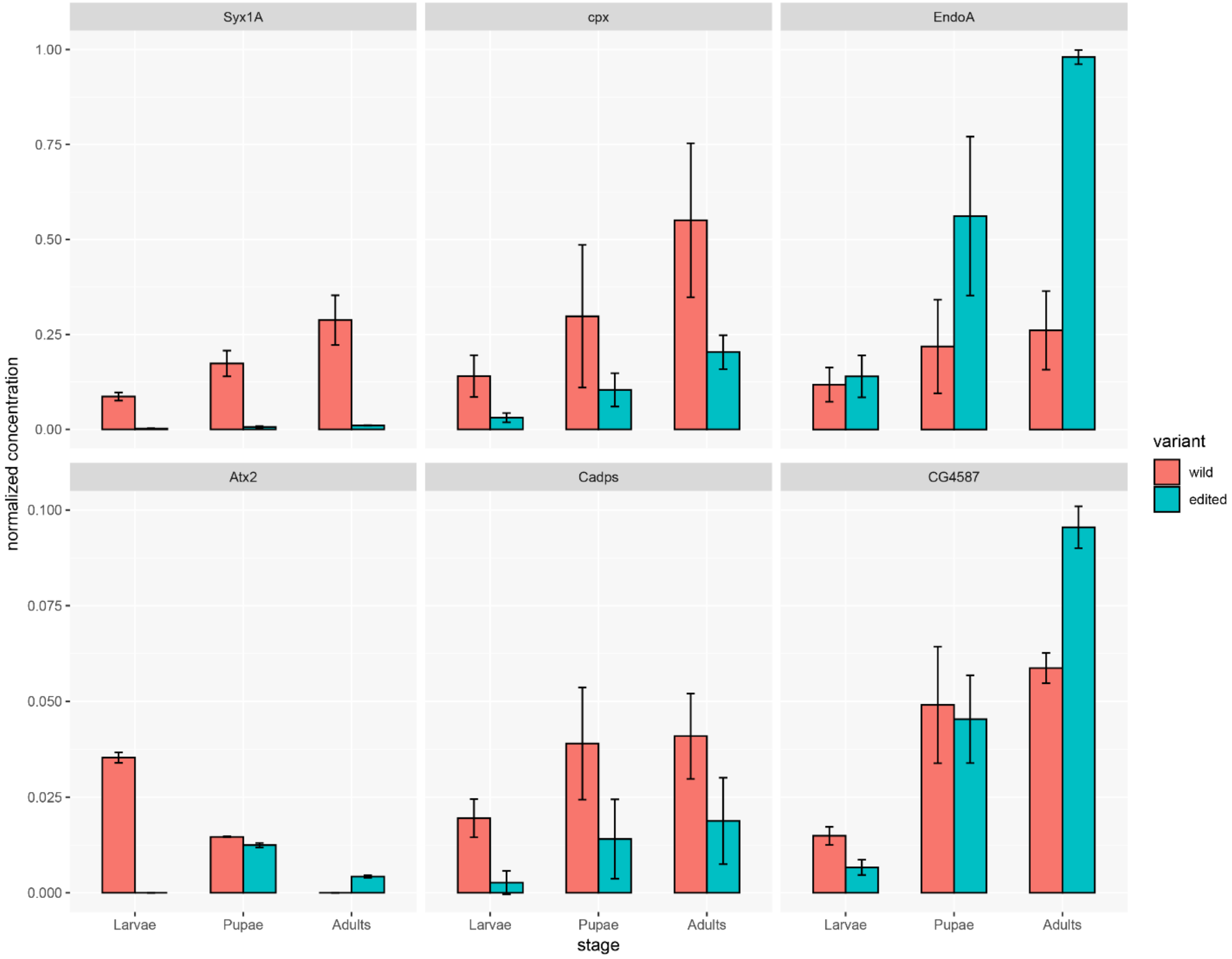


**Supplementary Figure 4. Concentrations of edited and unedited peptides normalized by total peptide content obtained for different biological replicates in the brain at life cycle phases of *D. melanogaster*.**

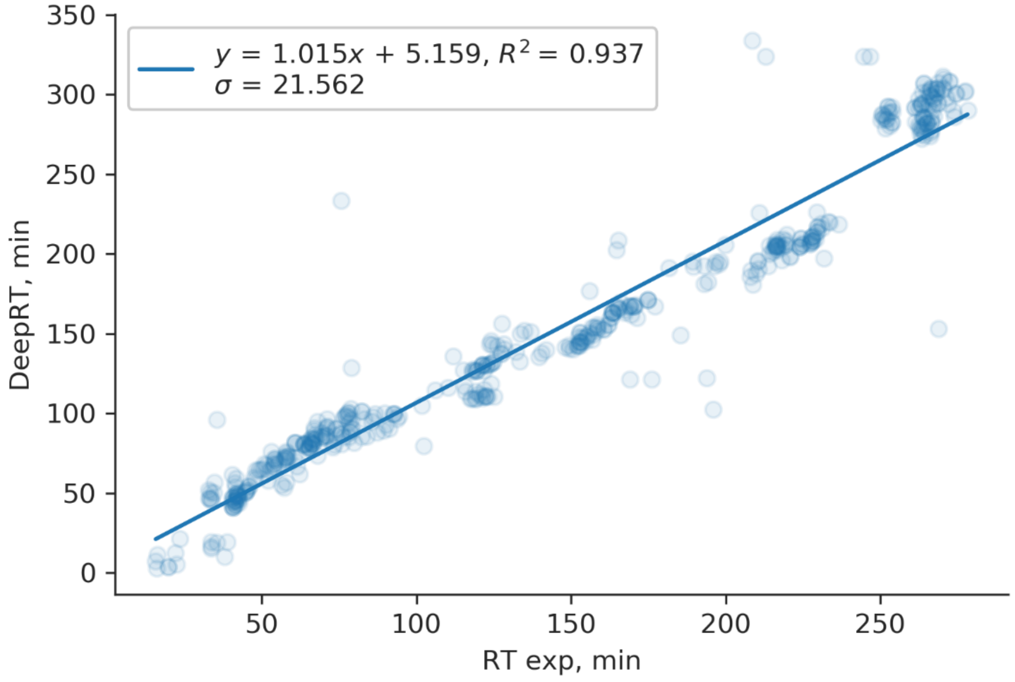


**Supplementary Figure 5. A linear plot representing relations between predicted and experimental retention times (RT) for each PSM.**
